# Assessing the role of long-noncoding RNA in nucleus accumbens in subjects with alcohol dependence

**DOI:** 10.1101/583203

**Authors:** Gowon O. McMichael, John Drake, Eric Sean Vornholt, Kellen Cresswell, Vernell Williamson, Chris Chatzinakos, Mohammed Mamdani, Siddharth Hariharan, Kenneth S. Kendler, Michael F. Miles, Gursharan Kalsi, Brien P. Riley, Mikhail Dozmorov, Silviu-Alin Bacanu, Vladimir I. Vladimirov

## Abstract

Recently, long noncoding RNA (lncRNA) were implicated in the etiology of alcohol dependence (AD). As lncRNA provide another layer of complexity to the transcriptome, assessing their expression in the brain is the first critical step towards understanding lncRNA functions in AD. To that end, we profiled the expression of lncRNA and protein coding genes (PCG) in nucleus accumbens (NAc) from 41 subjects with AD and 41 controls. At false discovery rate (FDR) of 5%, we identified 69 and 309 differentially expressed lncRNA and PCG genes, respectively. Using co-expression network analyses, we identified three lncRNA and five PCG modules significantly correlated with AD at Bonferroni adj. p≤0.05. To better understand lncRNA functions, we integrated the lncRNA and PCG hubs from the significant AD modules; at FDR of 5%, we identified 3 151 positive and 2 255 negative correlations supporting the functional role of lncRNA in the development of AD. Gene enrichment analysis revealed that PCG significantly correlated with lncRNA are, among others, enriched for neuronal and immune related processes. To highlight the mechanisms, by which genetic variants contribute to AD, we integrated lncRNA and PCG hubs with genome-wide SNP data. At FDR≤0.3, we identified 276 expression quantitative trait loci (eQTL), affecting the expression of 20 and 256 lncRNA and PCG hubs, respectively. Our study is the first to profile lncRNA expression in nucleus accumbens in a large postmortem alcohol brain sample and our results may provide novel insights into the regulation of the brain transcriptome across disease.

## Introduction

Alcohol Use Disorder (AUD) is a chronic and debilitating disease with an estimated heritability around 50%(1). Previous postmortem brain expression studies have shown that chronic alcohol consumption leads to broad transcriptional changes in different brain regions(2;3). Gene expression studies in prefrontal cortex (PFC), have identified genes encoding to GABA_A_ receptor subunits or related to mitochondrial function(4;5) as well as in functions related to myelination, cell cycling, oxidative stress, and transcription(2;6–11). Expression studies in nucleus accumbens (NAc) and ventral tegmental area (VTA), have also revealed gene expression changes related to cell architecture, signaling, vesicle formation and synaptic transmission(7). These findings suggest that there are region-specific susceptibilities and adaptations to chronic alcohol consumption in the brain that are likely to have a distinct effect on the behavioral phenotypes comprising AUD(12).

Evaluation of the regulatory mechanisms underlying genetic differentiation is necessary to better understand the neurobiology of alcohol addiction(13). The recent emergence of long noncoding RNA (lncRNA) has provided an additional layer of transcriptional and translational control that could highlight important neurobiological mechanisms underlying AUD that could be missed if only the protein coding genome was studied. LncRNA are longer than 200 base pairs (bp) with limited or no protein coding potential(14) and while they have not been functionally fully characterized yet, they were shown to participate in chromatin remodeling(15), transcriptional and post-transcriptional regulation(16), and in sequestering miRNA(17;18). LncRNA have also been implicated in the plasticity of neuronal circuity(19) and in neurodegenerative and neuropsychiatric disorders including substance abuse and alcohol dependence(20;21); the potential role of lncRNA in the etiology of AUD was also further supported by several recent alcohol-related genome-wide association scans (GWAS)(22;23). Due to the complex nature of lncRNA functions, the relationship between lncRNA and alcohol consumption has been understudied; while miRNA roles in the control of gene expression and functions in the brains of subjects with AUD have been reported(2;24–27), to our knowledge the interactions between lncRNA and protein coding genes (PCG) are still largely undescribed.

Gene expression alone cannot explain the complex etiology of AUD and assessing lncRNA and PCG expression in the context of available genetic data is necessary to discern the genetic basis of AUD susceptibility. This approach models associations between genetic variants and gene expression as quantitative traits, i.e. expression quantitative trait loci (eQTL)(28–30). EQTLs can help discover unknown AUD risk loci or offer specific, testable, hypotheses for the genetic impact of polymorphisms associated with AUD(31–33). Linkage disequilibrium (LD) of eQTLs with genetic variants implicated in AUD can further establish a biological mechanism for disease-associated variants with no apparent functions; there is empirical evidence suggesting that eQTLs are over-represented among GWAS signals(34;35).

We hypothesize that lncRNA are involved in the neuropathology of AUD and that, on a molecular level, this involvement is manifested by differential patterns of expression between cases with alcohol dependence (AD) and controls. To better understand the lncRNA disease functions in subjects with AD we further performed a weighted gene co-expression network analysis (WGCNA) that led to identification of lncRNA and PCG modules significantly correlated with AD. Key genes from these lncRNA and PCG modules (i.e., ‘hubs’) were then correlated with each other to identify a set of interacting lncRNA and PCG modules. Since lncRNA are not yet annotated, we used system approaches to assign biological functions for the PCG hubs interacting with the lncRNA hubs. By integrating our genome-wide SNP and expression data, we also identified expression quantitative trait loci (eQTL) affecting the expression of lncRNA and PCG in NAc. In this study, our overall goal is to profile lncRNA and PCG expressions in NAc and test whether these are under the control of specific genetic elements. To the best of our knowledge this is the only study to comprehensively assess the lncRNA expression between subjects with AD and controls in postmortem AD case and control brain tissues.

## Methods and Materials

### Postmortem tissue

Brain tissue from 41 cases with AD and 41 controls was received from the Australian Brain Donor Program, New South Wales Tissue Resource Centre, which is supported by The University of Sydney, National Health and Medical Research Council of Australia, Schizophrenia Research Institute, National Institute of Alcohol Abuse and Alcoholism, and the New South Wales Department of Health (http://sydney.edu.au/medicine/pathology/trc/). (Suppl. Methods; Suppl. Table 1).

### RNA isolation and sample selection

Total RNA was isolated from 50mg of frozen tissue from NAc using the mirVana-PARIS kit (Thermo Fisher, Carlsbad, CA), following manufacturer’s protocols. RNA concentration was measured using the Quant-iT Broad Range RNA Assay kit (Life Technologies), and the RNA Integrity Number (RIN) was measured on the Agilent 2100 Bioanalyzer (Agilent Technologies, Inc., Santa Clara, CA).

### Expression arrays and data normalization

The RNA samples were assayed using the Arraystar Human lncRNA Array v3.0 (Rockville, MD, USA) that is designed to profile both lncRNA and PCG. The expression values were calculated and normalized using the 75^th^ percentile value for each microarray in the Agilent Feature Extraction text file, followed by Log_2_ transformation and batch effect removal in the Partek^®^ Genomics Suite^®^ (PGS) software, v6.6 (St. Louis, MO, USA). Probes with expression values outside the detectable ‘spike-in’ range in more than 50% of the sample, or if they were flagged as “*gIsWellAboveBG=0*” and “*gIsPosAndSignif=0*” in more than 75% of the arrays were excluded from the downstream analyses. Upon removal of low expressed probes, the final number of probes that were used in the subsequent analyses was 19,458. The efficiency of batch effect removal and overall array quality were assessed using principal components analysis (PCA) on the expression values in which each array is plotted along the first three principle components (PCs) to identify potential outliers. Of the 82 samples, 9 samples were excluded due to low RINs (i.e. RINs≤4), leaving 73 samples for the microarray run. Of the 73 samples run on the array, 8 samples (5 cases and 3 controls) did not load on two of the first three PCs and were removed from the subsequent analysis, leaving a final sample size of N=65.

## Statistical Analyses

### Gene Expression Analysis

The univariate gene expression analyses were performed in the Number Cruncher Statistical Software (NCSS) v11, using a robust multiple regression model(2). Prior to the main analyses, technical (e.g. RIN, PMI, RI) and biological (e.g. smoking, medication, age) confounds and the first 5 principle components (PCs) were used as covariates in the regression analysis. Microarray reliability was validated by assessing the expression of 5 genes at the Arraystar facilities using quantitative PCR (qPCR). The assessed genes showed very high correlation between the two platforms (Suppl. Methods and Suppl. Fig. 1).

### Weighted Gene Co-expression Network Analysis (WGCNA)

Gene co-expression networks were constructed separately for the lncRNA and PCG using the WGCNA v1.36 package in R environment (v3.5.2). While, WGCNA allows for the inclusion of additional variables, it does not correct for their effects, therefore the gene networks were built using the residuals from the regression model. WGCNA relies on pair-wise Pearson correlations to generate a signed similarity matrix, selecting for positive correlations only. The signed similarity matrix of the lncRNA and PCG expression data was raised to the lowest power (β=8 and β=11, respectively) that approximated a scale-free network topology (R^2^>0.90), to generate an adjacency matrix. A topological overlap measure (TOM) was calculated to assess transcript interconnectedness. A dissimilarity measure was calculated from the TOM and was subsequently used for average linkage hierarchical clustering. Module definition parameters included a minimum module size of 35 genes, *DeepSplit 4*, and a minimum module merge height of 0.8.

Following module definition, the first principal component of each module – the module eigengene (ME) – was calculated as a synthetic gene representing the expression profile of all genes within a given module. Modules are named by a conventional color scheme and then correlated to AD case-status, matching demographics and relevant covariates. Statistical significance was assessed at Bonferroni-adj. p≤ 0.05 (corrected for number of tested modules).

To identify the set of genes with a high module membership and phenotype association, (i.e. the hub genes), we applied two selection criteria: 1) genes with the high intra-modular connectivity (r^2^ ≥0.7) and 2) significantly correlated with AD(36).

### Gene Set Enrichment Analysis

Gene set enrichment analysis (GSEA) was used to detect known biological processes and pathways enriched within the PCG modules using GSEA v2.2.3 software from the Broad Institute as previously described(37;38). Individual gene lists for each of the PCG modules significantly correlated with AD were generated by rank-ordering all differentially expressed PCG by their module membership (MM) to each of the AD-associated modules. Prior to running GSEA, the transcript IDs were converted to HUGO Gene Nomenclature Committee (HGNC)(39). We derived the a *priori* gene sets from the Molecular Signatures Database v6.2 (MSigDB; http://www.broadinstitute.org/gsea/msigdb) from the Broad Institute. A total of 1329 gene sets from the Canonical Pathways subset of the C2: Curated Pathways collection of MSigDB were assessed. Default parameters were then applied to give a minimum and maximum *a priori* gene set size between 15 and 500 genes, respectively. The identified enriched pathways were further adjusted for multiple testing at FDR of 25%.

### Correlation Analyses

The expression values of lncRNA and PCG hubs from the significant modules, were correlated (Pearson r) with each and adjusted at FDR of 5%, separately for the negative and positive correlations. To better understand the biological significance of these correlations, we performed gene enrichment analysis, in the Co-lncRNA webtool (http://bio-bigdata.hrbmu.edu.cn/Co-LncRNA/) using KEGG pathways. The significance of the enrichment analysis was assessed by hypergeometric test and further adjusted for multiple testing at FDR of 10%(40).

### eQTL Detection

The genotype calls were generated as part of a larger meta-GWAS study(22) and were integrated with the lncRNA and PCG hubs expressions to identify expression quantitative trait loci (eQTLs) for the lncRNA and PCG. Following a previously described approach and to retain power only local, cis-eQTLs, were assessed(2). Specifically, SNPs located 1 mega base pairs (Mbp) upstream and downstream from the selected lncRNA and PCG hubs were filtered with Plink v1.07 to exclude variants in LD (R^2^≥ 0.7)(41). To reliably estimate the eQTL effects in a sample size of 65, we further considered only SNPs with a minor allele frequency (MAF) ≥10%. After LD pruning we retained 58 317 SNPs which were tested as eQTLs. SNP impact on gene expression was detected by MatrixEQTL software package in R in using a linear regression framework and adjusting for the potential effects of the first 5 PCs(42). The significant eQTL results were adjusted for multiple testing at FDR of q≤0.3.

### Test for GWAS association signals enrichment

The assessment of enrichment for low p-values was performed by using the SST test(43). Briefly, for the analysis we used the latest data from the Psychiatric Genomic Consortium (PGC) GWAS of AUD(44). As the postmortem brain sample is composed of subjects with European (EU) ancestry, for our analysis we used the EU sample of 46K subjects (Suppl. Methods).

## Results

### Chronic alcohol consumption leads to generalized gene expression changes between cases and controls

At the nominal p≤0.05 we identified a total of 2,859 probes differentially expressed between AD cases and controls, representing 676 lncRNA and 2,135 PCG that compromise 3.4% and 11% of the total gene pool assayed, respectively. Similar to previous studies, the overall number of differentially expressed genes was greater than expected by chance (hypergeometric p=1.3×10^−5^) suggesting that chronic alcohol intake leads to generalized changes in the brain transcriptome(2;45). At FDR of 5% we identified 69 lncRNA and 309 PCG. Interestingly among these we also identified a few pseudogenes with evidence for differential expression. An unsupervised hierarchical clustering of the standardized expression values (i.e., values shift to a mean of 0 and standard deviation of 1) of all differentially expressed PCG and lncRNA genes at the nominal p≤0.05 showed general patterns of association between gene expression and diagnosis (Suppl. Fig.2A and 2B). The significant results from the univariate gene expression analyses are provided in supplementary table (Suppl. Table 2).

### lncRNA and PCG show a disease relevant network pattern

While the gene expression analysis identified differentially expressed genes in NAc of subjects with AD, these do not provide an integrative view of the interaction between the lncRNA and mRNA genes. Therefore, we performed a weighted gene co-expression network analysis (WGCNA)(46;47) separately on the nominally expressed lncRNA and PCG at p≤0.05. We chose this threshold in order to: **i**) include genes with smaller effect size (which otherwise would be excluded with more stringent statistical criteria), **ii**) enrich for genes likely to play role in AD, and **iii**) retain a sufficient number of genes for building of gene co-expression networks. In the lncRNA network analysis, we identified a total of 5 modules (Fig. 1)(46), of which three modules (*M*_*brown*_, *M*_*blue*_, and *M*_*grey*_) were significantly correlated with AD at a Bonferroni adj. p≤0.01. In the PCG network analysis, we identified 11 modules, five of which were significantly correlated with AD at a Bonferroni adj. p≤0.005 (Fig.2). In addition to the main dichotomous case/control status, we also tested additional quantitative alcohol related phenotypes such as age of initiation of alcohol drinking, amount of daily alcohol consumption and years of drinking. We identified one module (*M*_*green*_) and two modules (*M*_*brown*_ and *M*_*green*_) as significantly correlated with amount of daily alcohol consumption at a Bonferroni adj. p≤0.05 in the lncRNA and PCG analyses, respectively. Interestingly, two of the five PCG modules were significantly correlated with both AD and amount of daily alcohol consumption, while in the lncRNA analysis, we did not observe overlap between modules associated with AD (i.e. *M*_*brown*_ and *M*_*blue*_) and daily alcohol consumption (*M*_*green*_). Full tables with module size, correlations and p-values for all PCG and lncRNA modules correlated with AD and daily alcohol consumption are provided in supplementary tables (Suppl. Tables 3 and 4).

**Fig. 1.**
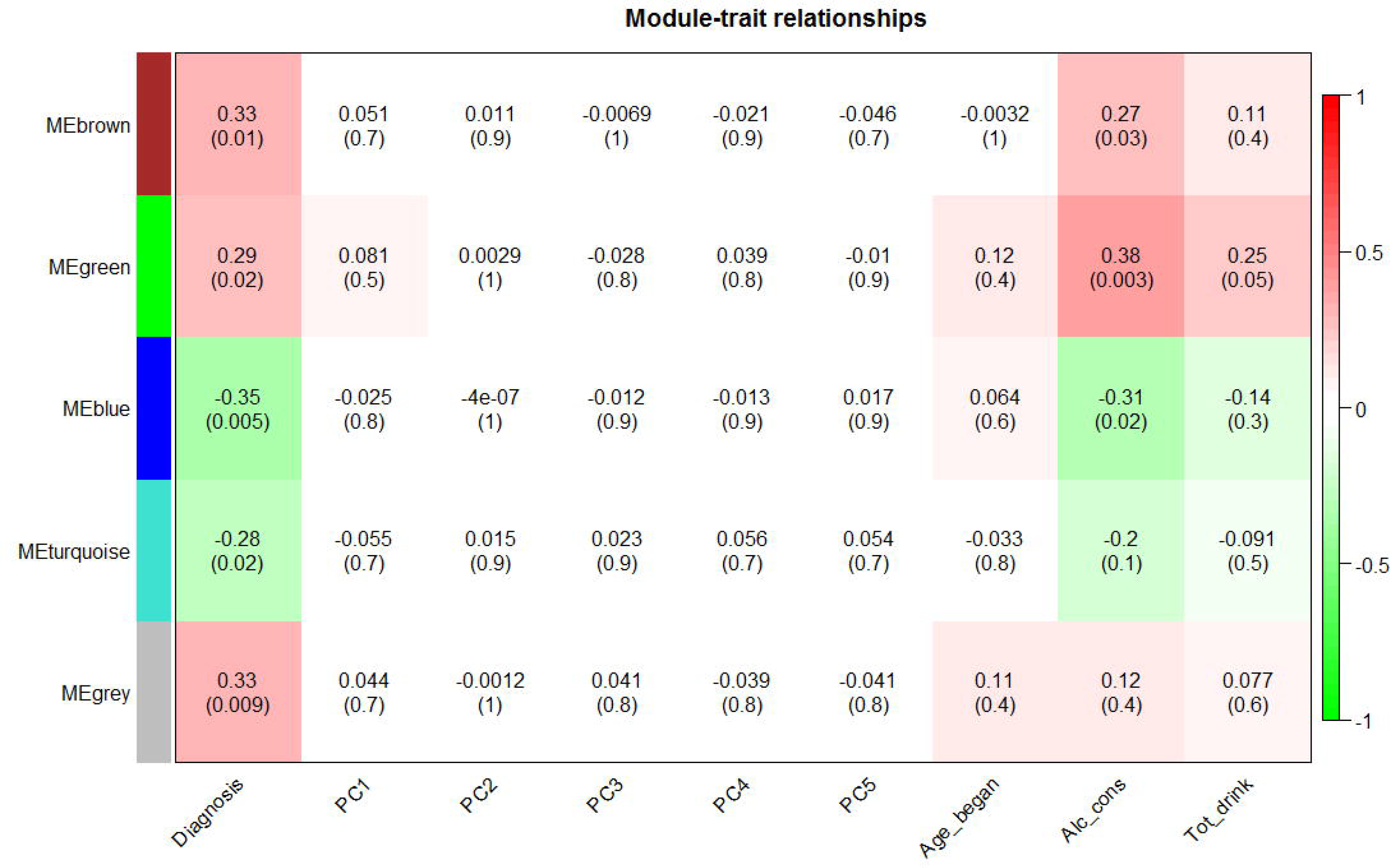
Module-trait relationships for lncRNA. The residuals of the expression values used to generate the lncRNA module MEs are correlated (Pearson) to the dichotomous AD case/control status (Diagnosis) and to quantitative alcohol measures such as daily alcohol consumptions (Alc-Cons), total amount of drinks (Tot_drinks) and initial age of drinking (Age_began). The lncRNA modules were also correlated to the first 5 PCs to assess for confounding. P-values shown are unadjusted for multiple testing. After adjusting for number of modules tested, ME_*brown*_, ME_*blue*_, and ME_*grey*_, are significantly correlated with AD case-status (Class) and ME_*green*_ with daily alcohol consumption.

**Fig. 2.**
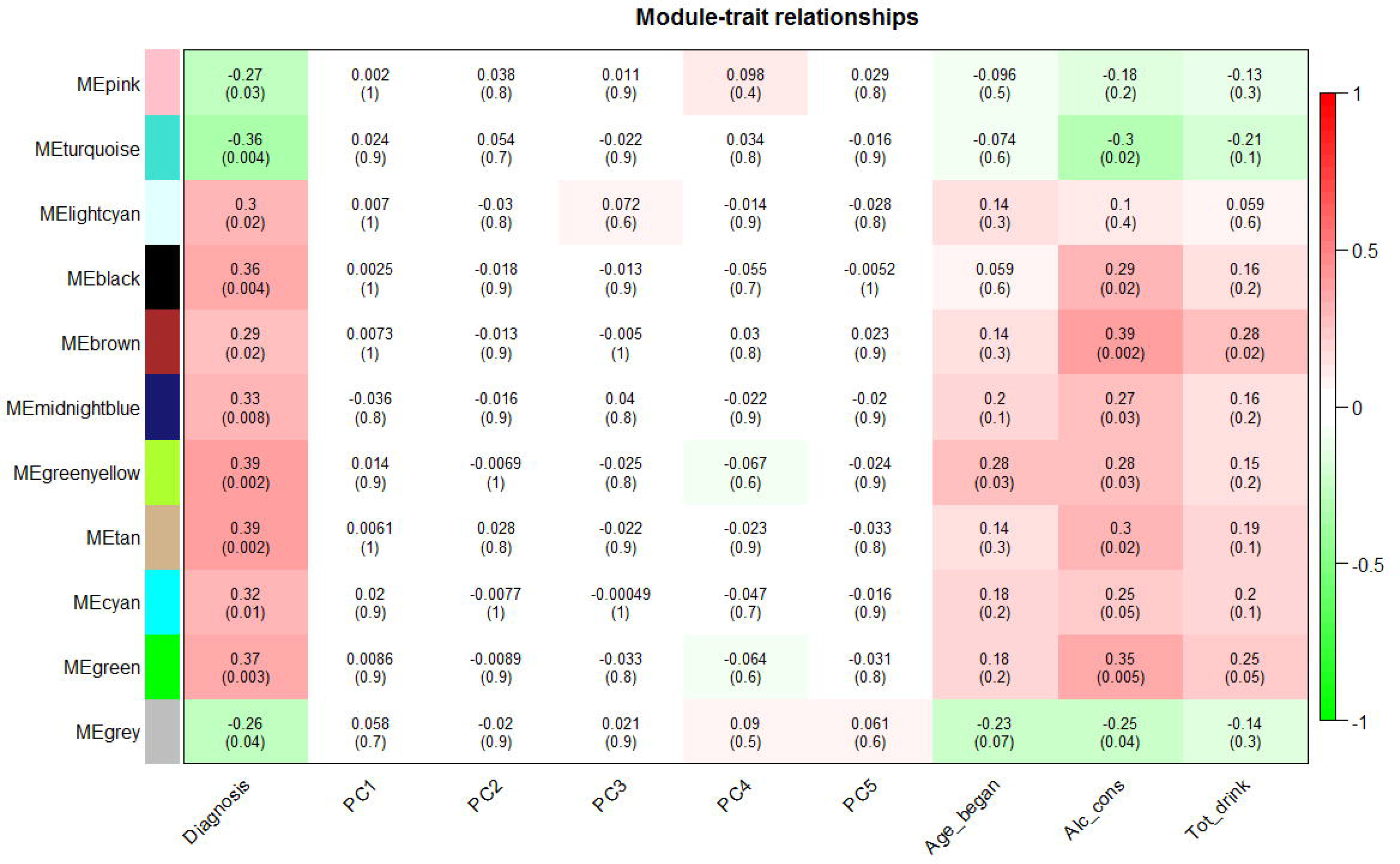
Module-trait relationships for protein coding genes (PCG). The PCG module MEs are correlated (Pearson) to the dichotomous AD case/control status (Diagnosis) and to quantitative alcohol measures such as daily alcohol consumptions (Alc-Cons), total amount of drinks (Tot_drinks) and initial age of drinking (Age_began). The lncRNA modules were also correlated to the first 5 PCs to assess for confounding. P-values shown are unadjusted for multiple testing. After adjusting for number of modules tested, ME_*turquoise*_, ME_*black*_, ME_*greenyellow*_, ME_*tan*_ and ME_*green*_ are significantly correlated with AD case-status (Class) and M_*brown*_ and ME_*green*_ also with daily alcohol consumption.

To ensure the robustness of our gene co-expression networks and reduce the potential influence of outlier samples on network structure, we used the robust ‘*boostrapped*’ version of WGCNA (rWGNCA). We performed 100 iterations in which networks were created after randomly subsetting 2/3 of the total samples as previously suggested(48). The resulting 100 networks were then merged into one large, final consensus network; the individual sub-networks showed reasonably high consistency with the final lncRNA and mRNA networks (Suppl. Fig.3A and 3B).

In a scale-free network topology, ‘hubs’ are the most highly connected genes (of which there are relatively few among all the nodes within a network); thus, we also identified the hub genes for the significant modules. Of the three lncRNA modules associated with AD, M_*blue*_ was the largest containing 200 genes, M_*brown*_ contained 105 genes and M_*grey*_ 73 genes. As the genes in M_*grey*_ are unassigned we attempted to identify hubs only in M_*blue*_ and M_*brown*_. Of the 305 transcripts clustered in the two modules, 70 transcripts were considered as candidate hubs, while among the five PCG modules we identified a larger number of hubs (N=368).

### AD gene modules are enriched in alcohol related processes

Previous studies have shown that co-expressed genes are enriched for biologically relevant functions [49-51]. We used a gene set enrichment analysis (GSEA) to assess for enrichment of cellular process and biological functional categories in modules associated with AD. The gene lists for each module were generated by ranking all 2185 differentially expressed transcripts according to their MM to each of the five significant modules as previously described(49). Using the default parameters in GSEA, at FDR of 25%, we identified 38 *a priori* gene sets significantly enriched in four of the five PCG modules associated with AD (Table 1). Among these, M_*turquoise*_ and M_*greenyellow*_, showed the highest and M_*green*_ the lowest numbers of enriched GO terms, respectively. The 4 modules showed distinct patterns of enrichment, with *M*_*turquoise*_ enriched for gene sets involved in neurodegenerative disorders (i.e., Alzheimer, Parkinsons and Huntingtons diseases) as well as gene sets enriched for neuronal system, neurotrophins signaling and oxidative stress, while *M*_*tan*_ and *M*_*greenyellow*_ were enriched for immune related processes and for genes involved in metabolic processes among others.

**Table 1.**

| Name | Module Color | SIZE | ES | NES | NOM p-val | FDR q-val |
| --- | --- | --- | --- | --- | --- | --- |
| REACTOME_GENERIC_TRANSCRIPTION_PATHWAY | Green | 18 | 0.39174604 | 2.0209973 | 0.00617284 | 0.09609686 |
| NABA_MATRISOME | Greenyellow | 52 | 0.29716566 | 2.5383055 | 0 | 0.002136364 |
| REACTOME_GENERIC_TRANSCRIPTION_PATHWAY | Greenyellow | 18 | 0.44453996 | 2.271343 | 0.003868472 | 0.011002417 |
| REACTOME_CYTOKINE_SIGNALING_IN_IMMUNE_SYSTEM | Greenyellow | 16 | 0.36958155 | 1.8165785 | 0.01814516 | 0.052214585 |
| REACTOME_TRANSMEMBRANE_TRANSPORT_OF_SMALL_MOLECULES | Greenyellow | 24 | 0.30484733 | 1.7894949 | 0.009727626 | 0.055684347 |
| KEGG_FOCAL_ADHESION | Greenyellow | 25 | 0.28989616 | 1.7495718 | 0.031809144 | 0.056554362 |
| NABA_MATRISOME_ASSOCIATED | Greenyellow | 32 | 0.263354 | 1.7666267 | 0.026476579 | 0.057339538 |
| NABA_CORE_MATRISOME | Greenyellow | 20 | 0.33983636 | 1.8274673 | 0.011538462 | 0.058284443 |
| KEGG_LEUKOCYTE_TRANSENDOTHELIAL_MIGRATION | Greenyellow | 15 | 0.39268157 | 1.8297838 | 0.017045455 | 0.06925771 |
| REACTOME_FATTY_ACID_TRIACYLGLYCEROL_AND_KETONE_BODY_METABOLISM | Greenyellow | 16 | 0.39104077 | 1.8677335 | 0.01417004 | 0.07026 |
| REACTOME_METABOLISM_OF_LIPIDS_AND_LIPOPROTEINS | Greenyellow | 40 | 0.25898334 | 1.8926 | 0.009784736 | 0.084817424 |
| REACTOME_CYTOKINE_SIGNALING_IN_IMMUNE_SYSTEM | Tan | 30 | 0.5582678 | 3.6189888 | 0 | 0 |
| REACTOME_INTERFERON_SIGNALING | Tan | 23 | 0.6121739 | 3.4482594 | 0 | 0 |
| REACTOME_IMMUNE_SYSTEM | Tan | 80 | 0.31643414 | 3.1712813 | 0 | 0 |
| REACTOME_INNATE_IMMUNE_SYSTEM | Tan | 20 | 0.51097137 | 2.762049 | 0 | 0 |
| REACTOME_GENERIC_TRANSCRIPTION_PATHWAY | Tan | 16 | 0.49438202 | 2.3938062 | 0 | 0.001711724 |
| NABA_MATRISOME | Tan | 52 | 0.27569824 | 2.2962203 | 0 | 0.002778108 |
| REACTOME_ANTIGEN_PROCESSING_CROSS_PRESENTATION | Tan | 15 | 0.46424875 | 2.1254985 | 0.001964637 | 0.007368235 |
| NABA_MATRISOME_ASSOCIATED | Tan | 36 | 0.28877163 | 2.0132291 | 0.008130081 | 0.014373438 |
| REACTOME_ADAPTIVE_IMMUNE_SYSTEM | Tan | 49 | 0.22688648 | 1.8398738 | 0.013861386 | 0.036948614 |
| REACTOME_HIV_INFECTION | Tan | 21 | 0.30654764 | 1.7100884 | 0.026871402 | 0.06993838 |
| REACTOME_CYTOKINE_SIGNALING_IN_IMMUNE_SYSTEM | Tan | 16 | 0.40826204 | 1.9591156 | 0.005813954 | 0.076536424 |
| REACTOME_HOST_INTERACTIONS_OF_HIV_FACTORS | Tan | 15 | 0.35319522 | 1.6167567 | 0.032 | 0.10237446 |
| NABA_ECM_REGULATORS | Tan | 15 | 0.3412781 | 1.5847824 | 0.04637097 | 0.10880603 |
| REACTOME_TCA_CYCLE_AND_RESPIRATORY_ELECTRON_TRANSPORT | Turquoise | 36 | 0.52024674 | 3.5809069 | 0 | 0 |
| REACTOME_RESPIRATORY_ELECTRON_TRANSPORT | Turquoise | 27 | 0.5581451 | 3.4332826 | 0 | 0 |
| REACTOME_RESPIRATORY_ELECTRON_TRANSPORT_ATP_SYNTHESIS_BY_CHEMIOSMOTIC_C | Turquoise | 31 | 0.51506513 | 3.377205 | 0 | 0 |
| KEGG_HUNTINGTONS_DISEASE | Turquoise | 43 | 0.43448094 | 3.345477 | 0 | 0 |
| KEGG_ALZHEIMERS_DISEASE | Turquoise | 38 | 0.4251922 | 3.100907 | 0 | 0 |
| KEGG_OXIDATIVE_PHOSPHORYLATION | Turquoise | 36 | 0.40677297 | 2.9108124 | 0 | 0 |
| KEGG_PARKINSONS_DISEASE | Turquoise | 36 | 0.40677297 | 2.8989124 | 0 | 0 |
| KEGG_CARDIAC_MUSCLE_CONTRACTION | Turquoise | 15 | 0.5186309 | 2.4297206 | 0 | 0.001646979 |
| REACTOME_NGF_SIGNALLING_VIA_TRKA_FROM_THE_PLASMA_MEMBRANE | Turquoise | 15 | 0.47475696 | 2.2123554 | 0.002136752 | 0.006105823 |
| REACTOME_NEURONAL_SYSTEM | Turquoise | 27 | 0.30835184 | 1.9067711 | 0.014522822 | 0.045417827 |
| REACTOME_METABOLISM_OF_MRNA | Turquoise | 17 | 0.365826 | 1.831359 | 0.007968128 | 0.062365737 |
| REACTOME_SIGNALLING_BY_NGF | Turquoise | 22 | 0.31402782 | 1.790927 | 0.02008032 | 0.07278288 |
| REACTOME_PLATELET_ACTIVATION_SIGNALING_AND_AGGREGATION | Turquoise | 17 | 0.35091397 | 1.7582463 | 0.013916501 | 0.081596814 |
| REACTOME_TRANSMISSION_ACROSS_CHEMICAL_SYNAPSES | Turquoise | 19 | 0.32222223 | 1.6900269 | 0.026262626 | 0.10649991 |

### lncRNA and PCG show a complex pattern of interactions

Since lncRNA can function as potential molecular “scaffolders”(50;51), i.e. by bringing different sets of genes near each other, we were interested to identify PCG hubs significantly correlated with lncRNA hubs. Limited studies have shown that lncRNA can affect gene functions by interfering with the expression of their specific gene targets(52). We integrated the lncRNA and PCG hubs by correlating the expression values of lncRNA and PCG hubs across the significant modules. At FDR of 5% we observed 5,404 correlations, of which the number of positive correlations (N=3,150) significantly exceeded the number of negative (N=2,254) correlations (Mann-Whitney U test p=1E-36). In these analyses, we observed generalized as well as module specific correlations. For example, hubs from the blue and brown lncRNA modules showed distinct either negative or positive correlations, respectively with hubs from the black PCG module (Fig.3A and 3B). Similarly, hubs from the blue lncRNA module were exclusively positively correlated with the hubs from M_*turqoise*_ PCG module, while hubs from the brown lncRNA module were negatively correlated with M_*turqoise*_. We further observed that the unique module correlations also drove the strongest positive and negative individual lncRNA/PCG hubs correlations. For example, the strongest negative correlation (Pearson r=-0.72, p=1.63E-11, q=2.07E-8) was observed between lncRNA (G075391) a hub gene in the brown lncRNA module on chromosome 7 and VPS51 (a member of the vacuolar protein sorting-associated protein 51 family) a hub from *M*_*turquoise*_ on chromosome 11. Further exploring the negative correlation between these two hubs we observed a moderate mediating impact of G075391 on the expression of VPS51 between cases and controls (ANCOVA, F=5.07, df=1, p=0.028). In the gene expression analysis, G075391 was overexpressed in the alcohol subjects, while VPS51 was underexpressed (Fig. 4) suggesting that one potential mechanism by which G075391 contributes to the neuropathology of AUD is by reducing the expression of VPS51. The entire list of positive and negative lncRNA/PCG hub correlations at FDR of 5% is provided in supplementary table (Suppl. Table 5).

**Fig. 3.**
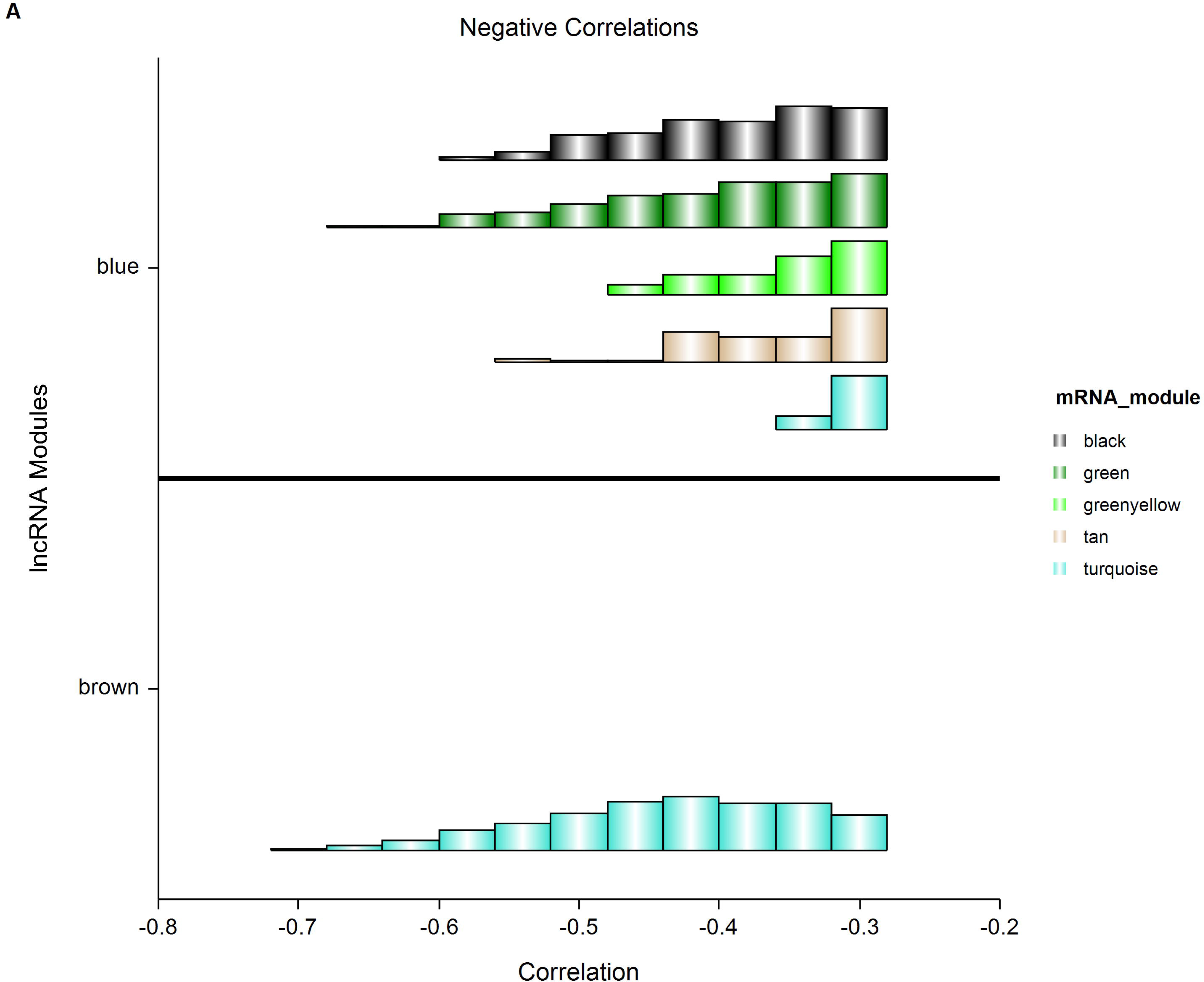
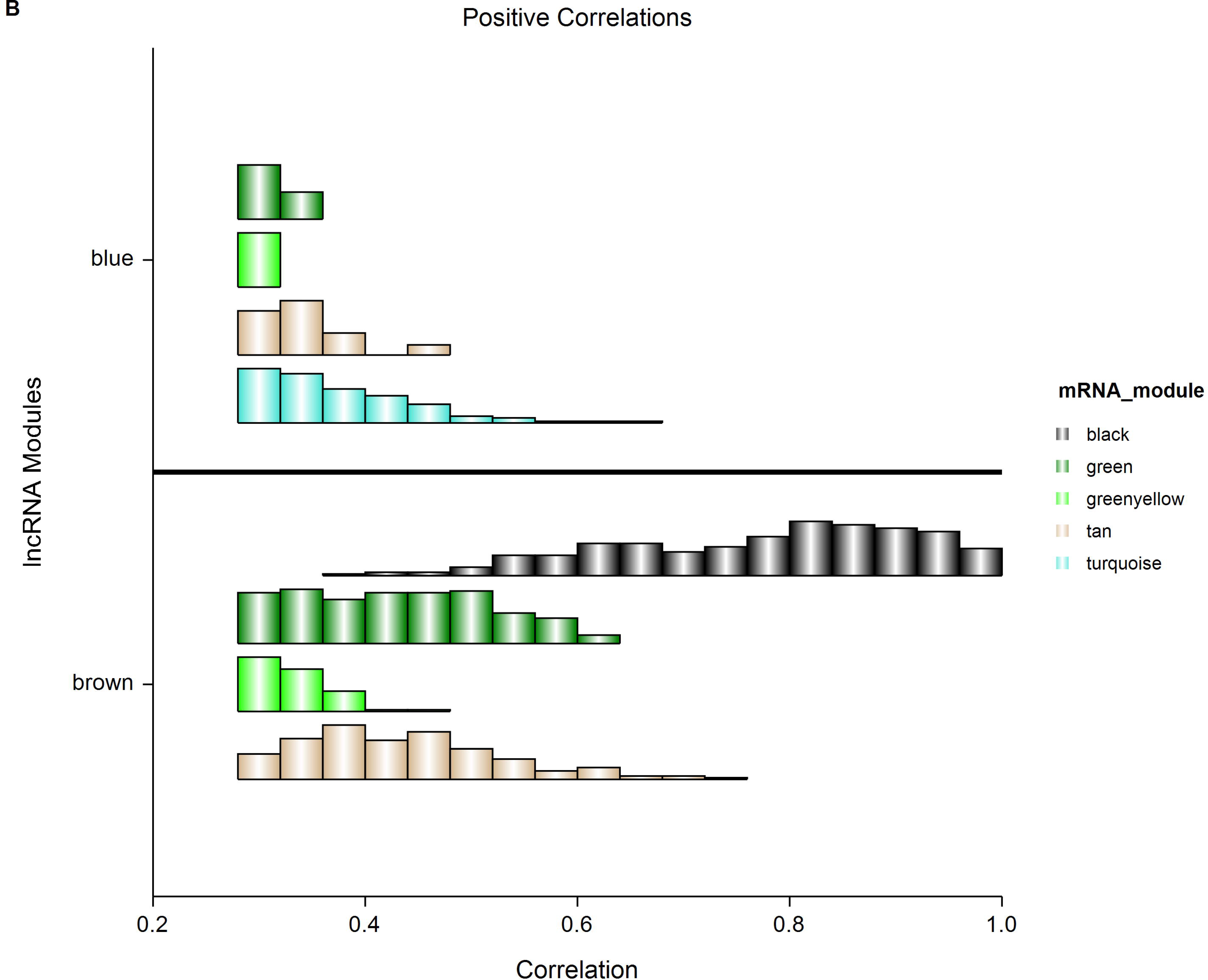
Histogram of lncRNA/PCG module correlations. (**A**) lncRNA ME_*brown*_ shows negative correlation with PCG ME_*turquoise*_, while lncRNA ME_*blue*_ showed negative correlations with all five significant PCG MEs. (**B**) PCG ME_*black*_ shows a positive correlation only with lncRNA ME_*brown*_ and PCG ME_*turquoise*_ shows a positive correlation only with lncRNA ME_*blue*_.

**Fig. 4.**
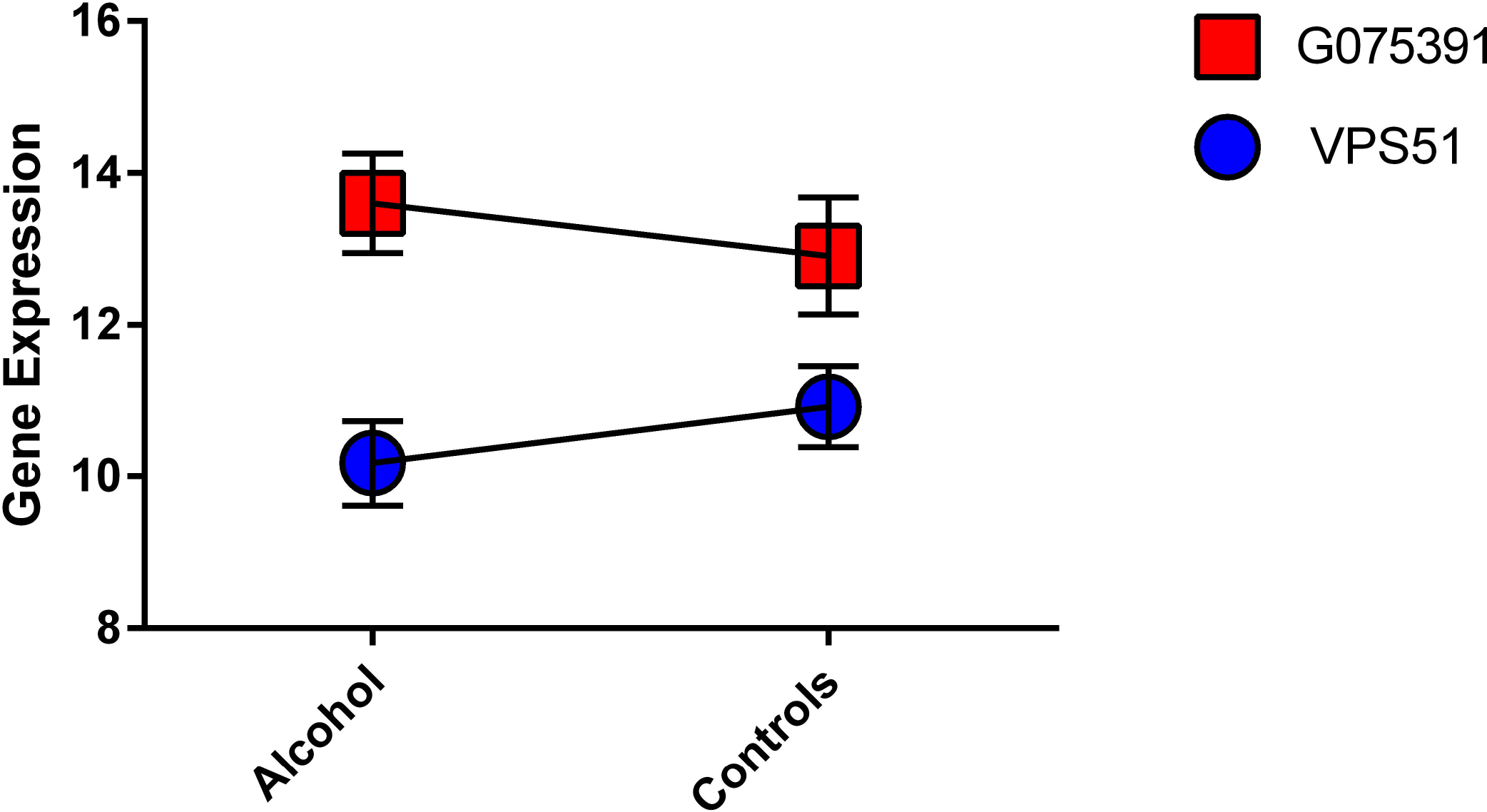
Plot of the mediating effect of lncRNA (G075391) on the expression of PCG Vacuolar Protein Sorting-Associated Protein 51 (VPS51) in cases and controls.

To better understand the overall functional role of the two significant lncRNA modules, we performed a pathway enrichment analyses using the Co-lncRNA webtool(40) for the negatively and positively correlated lncRNA and PCG modules. We observed distinct pathway enrichments, in which PCG modules negatively correlated with the lncRNA modules (*M*_*blue*_) were enriched for pathways related to the complement system, cell adhesion and toll-like receptors, while the positively correlated PCG modules were enriched for pathways belonging to neurodegenerative disorders such as Parkinson’s, Alzheimer and Huntington disease and in pathways involved in oxidative phosphorylation, long-term potentiation among others. PCG modules positively correlated with lncRNA M_*brown*_ module were enriched for lysosome trafficking, while PCG modules negatively correlated with lncRNA M_*brown*_ were enriched for genes involved in calcium signaling, long-term potentiation and neurotrophin signaling (Suppl Figs. 4A-4O). The PCG M_*brown*_ significantly correlated with daily alcohol consumption and negatively correlated with lncRNA M_*green*_ was enriched for gene pathways involved in focal adhesion, MAPK signaling, peroxisome (Suppl. Figs. 5A-5D).

### Genetic variants affect lncRNA and PCG expression in a disease specific manner

To understand the underlying genetic mechanisms of AD risk polymorphisms we further integrated the lncRNA and PCG hubs expression data from our significant modules with previously collected genotypic data(22). Due to the small sample size, we focused on testing cis-eQTLs (defined within 1Mbp up- and down-stream from the tested loci) only. At FDR of q≤0.3 we identified 293 eQTLs that impact hubs expression, with disproportionate number of eQTLs for the PCG (n=282) vs. lncRNA (n=11) hubs (Suppl. Table 6). In a follow up analysis, we further tested these 293 eQTLs for an interaction between genotype and disease status; at FDR of 5% we identified 31 interactions that mediate the expression of 30 PCG hubs, and one lncRNA hub (G054549) on Chr. 3 (Table 2). The most significant interaction we observed was between FKBP prolyl isomerase 5 (FKBP5) and rs9394312 on Chr. 6 (Fig.5).

**Fig. 5.**
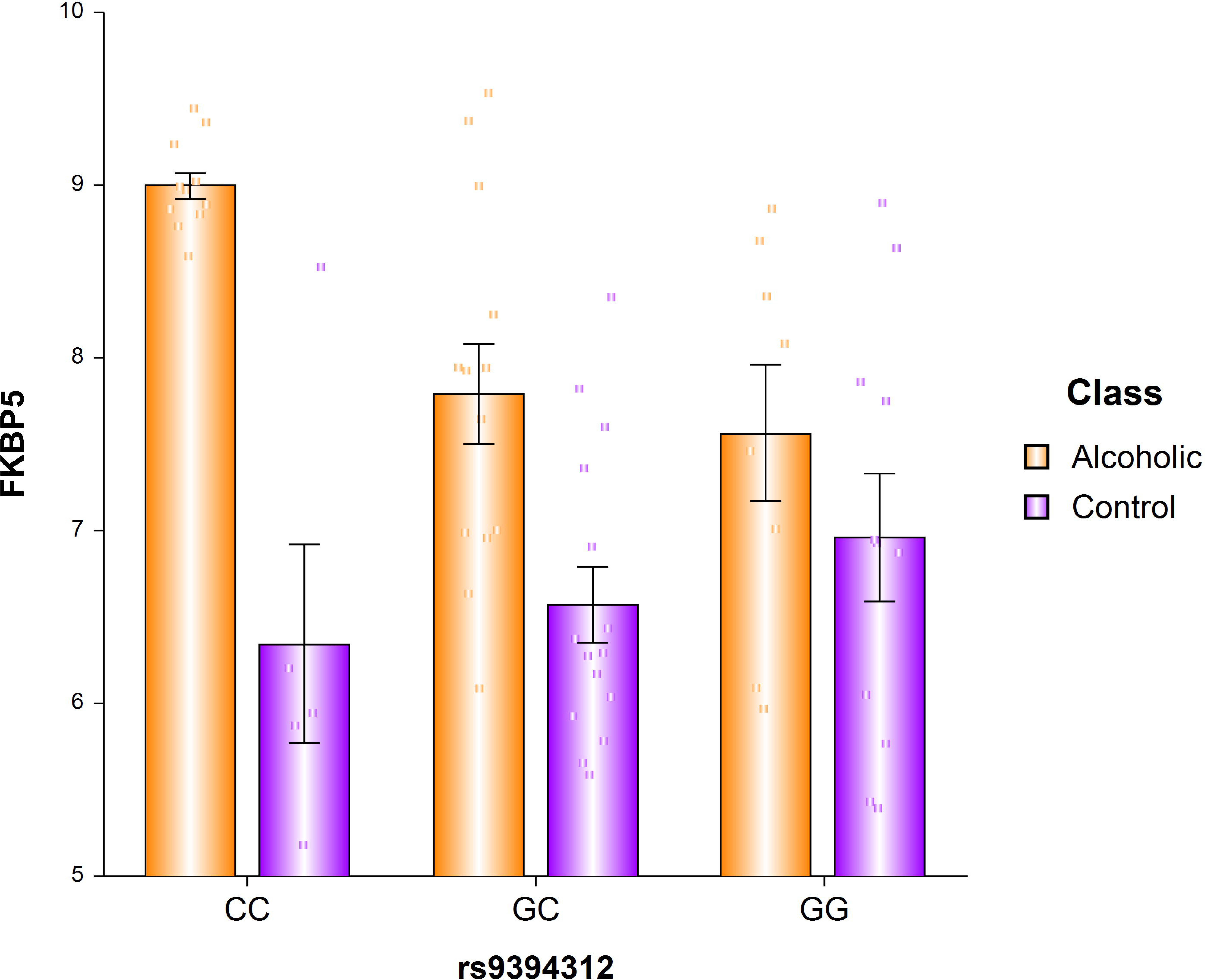
Plot showing a SNP interaction effect of disease status on the expression of FKBP5. The minor allele of rs9394312 confers a higher FKBP5 expression in controls and lower expression in cases.

**Table 2.** Statisitstics associated with SNP x Disease status interaction on gene expression

| Gene Symbol | transcript_type | b<sub>(i)</sub> | t-value | p-value | q-value |
| --- | --- | --- | --- | --- | --- |
| CP | protein_coding | -4339.099284 | -4.17296636 | 1.06E-04 | 0.0021200 |
| FKBP5 | protein_coding | -194.1447756 | -3.892794888 | 2.66E-04 | 0.0026600 |
| CFB | protein_coding | -816.8607338 | -3.575516617 | 7.29E-04 | 0.0048600 |
| CP | protein_coding | -2674.651616 | -3.216831651 | 0.002155381 | 0.0090439 |
| VWF | protein_coding | -10483.74203 | -3.200519733 | 0.002260962 | 0.0090439 |
| CP | protein_coding | -2902.166648 | -2.992404662 | 0.004111624 | 0.0118070 |
| SERPINB1 | protein_coding | -469.2666009 | -2.978236654 | 0.004278888 | 0.0118070 |
| MS4A6A | protein_coding | -892.6519535 | -2.941793914 | 0.004738668 | 0.0118070 |
| FKBP5 | protein_coding | -160.8875175 | -2.900623299 | 0.00531313 | 0.0118070 |
| PEF1 | protein_coding | 1051.251682 | 2.835312324 | 0.006358287 | 0.0123182 |
| FKBP5 | protein_coding | -161.1241294 | -2.808411694 | 0.006841617 | 0.0123182 |
| CFB | protein_coding | -852.5263206 | -2.779889041 | 0.007390944 | 0.0123182 |
| ETFB | protein_coding | 3725.712034 | 2.710880859 | 0.008892136 | 0.0136802 |
| CD14 | protein_coding | 4597.22194 | 2.662656152 | 0.010102021 | 0.0144315 |
| CFB | protein_coding | -455.8770002 | -2.60100512 | 0.011867568 | 0.0150922 |
| CP | protein_coding | -2998.120486 | -2.59435938 | 0.012073783 | 0.0150922 |
| CP | protein_coding | -2937.772237 | -2.556507693 | 0.01331174 | 0.0156609 |
| KIF19 | protein_coding | -176.2846059 | -2.478319252 | 0.016240449 | 0.0176072 |
| FKBP5 | protein_coding | -135.5325182 | -2.464543997 | 0.016812949 | 0.0176072 |
| G054549 | long noncoding | -76.58008707 | -2.44596443 | 0.01761392 | 0.0176072 |
| CFI | protein_coding | -479.7336288 | -2.422108455 | 0.018692651 | 0.0176072 |
| SLC2A5 | protein_coding | -670.9679148 | -2.407797313 | 0.019367912 | 0.0176072 |
| LIPA | protein_coding | 2571.97528 | 2.377762233 | 0.02085671 | 0.0181363 |
| CFB | protein_coding | -487.8252969 | -2.343161594 | 0.022698128 | 0.0189151 |
| IRF1 | protein_coding | -893.4275376 | -2.183954119 | 0.033170244 | 0.0259525 |
| C1QB | protein_coding | -794.4082368 | -2.16374536 | 0.034765474 | 0.0259525 |
| FKBP5 | protein_coding | -143.4456841 | -2.160400441 | 0.035035906 | 0.0259525 |
| CLIC1 | protein_coding | -8449.40885 | -1.9737918 | 0.038346681 | 0.0381048 |
| SMARCD3 | protein_coding | -4225.619536 | -1.917513507 | 0.040278624 | 0.0415715 |
| CLIC1 | protein_coding | -6404.73533 | -1.874201051 | 0.046122845 | 0.0440819 |
| CD14 | protein_coding | -3360.788427 | -1.857196165 | 0.048544994 | 0.0442226 |

### GWAS signals of AUD show suggestive enrichment in our lncRNA specific eQTLs

While we and others have shown alcohol related eQTLs are enriched for GWAS signals of AUD and other alcohol related phenotypes (ARPs) in the protein coding genome(53), no such studies have been performed to interrogate for such enrichment in lncRNA genes. To increase power of our test and be rather inclusive, we selected all eQTLs significant at the nominal p≤0.05. At this threshold level we have identified a total of 3064 eQTLs (366 lncRNA and 2698 PCG specific eQTLs). The assessment of enrichment was performed by using the SST test (Suppl. Text) and we detected only a suggestive enrichment (p=0.16) of AUD signals, most likely due to the underpowered alcohol PGC sample and our own postmortem brain sample.

## Discussion

The main goal of this study is to profile the expression patterns of the non-coding and coding transcriptome of NAc in a large sample of subjects with AD and controls. The NAc is a central component of the mesocorticolimbic system (MCL) and has been shown to be involved in addictive behaviors(54–56). By integrating GWAS data previously collected on the sample(22) with our expression data, we further attempted to reveal hitherto unknown relationships between lncRNA and PCG and to assess how this relationship is modulated by genetic variants in the brain. Unsurprisingly and in agreement with previous alcohol related postmortem brain expression studies, we identified differentially expressed PCG, suggesting that chronic alcohol consumption is associated with wide-spread transcriptomic changes in the brain. We further identified numerous lncRNA differentially expressed between alcoholic cases and control, supporting their involvement in the neuroadaptations associated with chronic alcohol consumption. Among the most significant differentially expressed PCG, were serpin peptidase inhibitor, clade A (alpha-1 antiproteinase, antitrypsin), member 3 (SERPINA3), interferon induced transmembrane protein 2 and 3 (IFITM2 and IFITM3) and solute carrier family 14 (urea transporter), member 1 (Kidd blood group) (SLC14A1), that were also identified from previous alcohol related postmortem brain gene expression studies(9;57). Interestingly, we also detected a few differentially expressed pseudogenes, such as the annexin A2 pseudogene 2 and 3 (ANXA2P2 and ANXA2P3) and interferon induced transmembrane protein 4 pseudogene (IFITM4P) on chromosomes 9, 10 and 6, respectively. Previous postmortem brain expression studies have shown ANXA2P2 to be involved in stress response pathways in chronic alcoholics(57), however, little is known about the functions of ANXA2P3 and IFITM4P; it appears that ANXA2P3 is involved in reducing lipids levels in response to statins(58) and in chronic obstructive pulmonary disease (COPD)(59), while no known functions were reported for IFITM4P.

A major limitation of the single gene expression analysis to identify and prioritize a set of genes associated with a given phenotype, is its inability to consider the complex molecular interactions between such identified genes(60); this is especially relevant in studying the non-coding transcriptome, which is poorly annotated, with many lncRNA genes still of unknown function. To address this limitation, we performed a weighted gene co-expression analysis (WGCNA), which allowed us to identify a set of co-regulated PCG and lncRNA. We identified three lncRNA modules and four PCG modules that were significantly correlated with AD. Among the significant lncRNA modules correlated with AD status was M_*grey*_. Genes in the grey module are frequently considered as ‘noise’, i.e. genes whose expression does not correlate with genes in the other modules, although there are suggestions that this interpretation may not always be accurate(61). In general, the genes in the grey module are usually ubiquitously expressed, exhibiting oscillating or highly variable patterns of expression; of course these could also be simply assigned to the wrong module(62). Regardless, one potential explanation for our observation could be that, at least some lncRNA, may impact the neuropathology of AD in isolation and not as a part of gene network. For example, other studies have also reported similar observation, where despite of their unassigned status, genes in M_*grey*_ show significant disease associations in certain phenotypes(63;64). Using the available quantitative measures of alcohol consumption, we further identified one lncRNA module and two PCG modules significantly correlated with amount of daily alcohol consumption after correction for multiple testing. Interestingly, while we observed same PCG modules to be significantly correlated with both AD status and quantitative alcohol phenotype, no such relationship was revealed for the lncRNA module, suggesting that different sets of lncRNA genes may impact these related, yet distinct, measures of alcohol related phenotypes. To further understand the potential regulatory functions of lncRNA, we performed a series of correlation analyses between the hub genes identified from the significant lncRNA and PCG modules. In these analyses we observed highly significant positive and negative correlations, in which the positive correlations significantly outnumbered the negative correlations; this is not entirely surprising, considering that the positive correlation can reflect both a genuine regulatory function as well as co-expression patterns between lncRNA and PCG. We further observed that the module specific correlations were driven exclusively by lncRNA belonging to individual modules, suggesting that individual modules are likely to contain a set of lncRNA with different and non-overlapping functions. Indeed, in our pathway enrichment analyses we observed the PCG modules negatively or positively correlated with the lncRNA modules are enriched for distinct and mostly non-overlapping biological processes, further corroborating the suggestion that lncRNA involved in the neuroadaptation to alcohol consumption likely have non-overlapping functions. From our GSEA analysis, we observed that similarly to other studies(2;45;65), some of the PCG modules are also enriched for immune related or neurodegenerative processes. Immune related responses to alcohol consumption in the brain has been well documented(66;67). Likewise, the neurodegenerative processes observed for some of our modules may reflect the nature of our postmortem sample composed exclusively from chronic alcoholics. This is not surprising considering that our sample is composed of subjects that have been exposed to alcohol for over two decades. Both animal and human genetic models have demonstrated that prolonged exposure to alcohol leads to activation of the microglial population with the concurrent activation of immune related processes in the brain(68;69). It has been well known, for over several decades now, that prolonged and excessive alcohol drinking leads to loss of a brain matter(70;71), and consequently activation of genes involved in neurodegenerative disorders.

Integrating our expression and genetic data led to identification of eQTLs that affect lncRNA and PCG expression. Having an integrated view on the expression of lncRNA and PCG we were also interested to identify eQTL that can impact both lncRNA and PCG expressions; we, however, failed to observe eQTLs that have a shared impact on lncRNA and PCG expressions. We further observed disproportionate numbers of eQTLs for PCG vs. lncRNA genes. This could reflect either a genuine biological mechanism, or conversely can be due to a lack of sufficient power to reliably identify eQTLs for lncRNA whose expression is associated with AD. Interestingly, among the eQTLs affecting PCG expression, certain polymorphisms showed a strong interaction effect between PCG expression and disease status. Among these, the strongest interaction effect was observed for FKBP5 and rs9394312. Several studies have implicated FKBP5 in the severity of alcohol withdrawal(72), alcohol drinking patterns in rodents(73), problematic drinking(74), as well as direct regulation of FKBP5 expression and phosphorylation by acute ethanol in mouse prefrontal cortex(75;76).

The importance of our postmortem brain expression study to assess lncRNA expression in nucleus accumbens is further supported by three recent alcohol related GWAS reporting genome-wide signals near lncRNA genes: i) a study conducted by Gelernter at al. (2014)(77) that implicated the ADH gene cluster on chromosome 4 and the LOC100507053 locus, ii) a study conducted by the COGA group(78) that identified a polymorphism near LOC151121 on chromosome 2, and iii) a study conducted by our own group that have reported several polymorphisms in LOC339975(22). However, in our study none of these three loci showed evidence for differential expression, which also agrees with our previous study reporting no evidence for differential expression of these loci, although we observed the expression of LOC339975 to be affected by the most significantly associated polymorphism (rs11726136) in NAc.

In summary, the main goal of this study was to test the hypothesis that lncRNA, as a novel class of none coding RNAs, may contribute to the neuropathology of alcohol dependence. In this study we identified a set of lncRNA and PCG differentially expressed between cases and controls. Our gene network analyses identified lncRNA and PCG modules significantly correlated with AD in NAc, which were shown to be enriched for immune related processes and neurodegeneration. We further identified distinct patterns of correlation between the lncRNA and PCG hubs with minimal overlap between modules, i.e. either positive or negative correlations, but not mixed. Gene pathway enrichment revealed that of genes whose functions were related to immune, neurological and neurodegenerative processes. Finally, we identified eQTLs that affect the expression of lncRNAs and PCG hubs, with some of these eQTLs showing a clear mediating effect on the PCG expression between cases and controls. Unlike previous studies though, here we saw only suggestive enrichment for association signals among the eQTLs, which could be attributed to either a limited importance of lncRNAs in the etiology of AD, or most likely due to our limited sample size.

While our study is novel in its approach to integrate genetic and molecular data in postmortem alcoholic brains as well as addressing important questions regarding lncRNA involvement in the etiology of AD, it is not without limitations. First, postmortem brain studies are observational as manipulation of living human subjects is not possible. Although the cross-sectional nature of these studies limits the causal inference we can make, we believe the eQTL analysis is a major step toward clarifying the directionality of these observations. Secondly, although our sample size (N=65) is by far the largest postmortem alcohol study to date, in comparison with other publically available postmortem brain expression samples(79) used to study neuropsychiatric disorders such as schizophrenia, bipolar disorder or major depression, is still prohibitively small. However, we believe that with our careful experimental design and implementation of integrative multivariate approaches, we can circumvent some of these limitations and further broaden our understanding of alcohol addiction processes. We hope that our study will provide additional molecular targets that will help translate these advances into effective therapeutic strategies for patients suffering with substance use disorders. Thus, in conclusion, we believe our study to be important in the field, as to the best of our knowledge, it is among the first such studies to address the role of lncRNA in the neuropathology of alcohol addiction paving the road for future such studies.

---

**Suppl. Fig. 1** Microarray expression data validation using quantitative PCR. Expression levels of five genes measured by the expression array-based approach were validated using quantitative PCR in all 65 postmortem Nucleus Accumbens (NAc) RNA samples.

**Suppl. Fig. 2** Unsupervised hierarchical clustering of the lncRNA and PCG differentially expressed at p≤0.05 was performed on the standardized expression values, which were centered at mean of 0 and SD of 1. The genes were clustered according to the similarity of their expression profile in PGS v.6.6 using complete linkage and Euclidean distance metrics. The color scheme of the Y-axis reflects the clustering based on diagnosis, while the color scheme of the X-axis is an arbitrary identification of the two large clusters.

**Suppl. Fig. 3** Robust, bootstrapped version of WGCNA (rWGCNA). The purpose of rWGCNA is to assess for potential influence of outlier samples on network structure. We performed 100 iterations in which networks were created after first randomly subsetting 2/3 of the total samples. The resulting 100 networks were then merged into one large, final consensus network. The individual sub-networks show consistent structure with each other and the final network for (**A**) lncRNA and (**B**) PCG.

**Suppl. Fig. 4** Figures show the enrichment of the PCG hubs positively and negatively correlated with lncRNA hubs in KEGG pathways for AD status. The pathways were derived from the Co-lncRNA web-tool(40). (A) PCG hubs from the ME_*green*_ positively correlated with lncRNA ME_*blue*_ are enriched for lysosome functions; (B) PCG hubs from the ME_*greenyellow*_ negatively correlated with lncRNA ME_*blue*_ are enriched for cell adhesions functions; (C) PCG hubs from the ME_*tan*_ negatively correlated with lncRNA ME_*blue*_ are enriched for genes with functions in complement system; (D) PCG hubs from the ME_*tan*_ negatively correlated with lncRNA ME_*blue*_ are also enriched for genes involved in Toll-like receptors; (E, F, G) PCG hubs from the ME_*turquoise*_ positively correlated with lncRNA ME_*blue*_ are enriched for genes involved in neurodegenerative diseases, i.e. Alzheimer’s, Huntington and Parkinson; (H, I, J, and K) PCG hubs from the ME_*turquoise*_ positively correlated with lncRNA ME_*blue*_ are also enriched for genes involved oxidative stress, glioma, long term potentiation, and spliceosome; (L) PCG hubs from the ME_*green*_ positively correlated with lncRNA ME_*brown*_ are enriched for genes involved in processes related to lysosome functions; (M, N, and O) PCG hubs from the ME_*turquoise*_ negatively correlated with lncRNA ME_*blue*_ are enriched for genes involved in calcium signaling, long term potentiation and neurotrophin signaling.

**Suppl. Fig. 5** Figures show the enrichment of the PCG hubs positively and negatively correlated with lncRNA hubs in KEGG pathways for daily alcohol consumption. (A, B, C, D) PCG hubs were enriched for processes related to focal adhesion, MAPK signaling, pathways in cancer, and peroxisome functions.

