## Supplementary material for "Assessing the role of long-noncoding RNA in nucleus accumbens in subjects with alcohol dependence": Suppl. Methods

**Supplementary Methods**

### 1. Postmortem Brain sample

Tissues from 41 AD cases and 41 controls were received from the Australian Brain Donor Program, New South Wales Tissue Resource Centre, which is supported by The University of Sydney, National Health and Medical Research Council of Australia, Schizophrenia Research Institute, National Institute of Alcohol Abuse and Alcoholism, and the New South Wales Department of Health (http://sydney.edu.au/medicine/pathology/trc/). Cases were excluded if they had an infectious disease (i.e. HIV/AIDS, hepatitis B or C, or Creutzfeldt-Jakob disease), an unsatisfactory agonal status (determined from the circumstances surrounding the death), post-mortem delays >48 hours or significant head injury. In addition to case status, age, sex, ethnicity, brain weight, brain pH, post-mortem interval (PMI), tissue hemisphere, clinical cause of death, blood toxicology at time of death, smoking status, neuropathology and liver pathology were also provided for each subject (Suppl. Table 1).

### 2. Array validation

The lncRNA arrays were validated at the Arraystar facilities by selecting five genes. The first-strand (reverse transcriptase (RT) reaction) cDNA was generated using SuperScript III Reverse Transcriptase (Invitrogen) following manufacturer's instructions. The reactions were performed in 25µl reaction volume and initially they heated to 65°C for 5 min, quenched on ice for 1 min, and kept at 50°C for 60 min. Following the RT reactions, the quantitative PCR reactions were performed in 10 µl volume reactions using the following parameters: initial denaturing for 10 min. at 95°C, followed by 40 amplification cycles of denaturing at 95°C for 10s and annealing/amplification at 60°C for 1 min. The PCR efficiency was assessed using a standard curve analysis with 10-fold serial dilutions (from 1 to $10^{-6}$).

The primer sets used to validate lncRNA results are listed in table 1

| Gene Name | Sequence (5' to 3') | Tm(°C) | Length of product(bp) |
|---|---|---|---|
| β-actin | Forward:5' GTGGCCGAGGACTTTGATTG3'<br>Reverse:5' CCTGTAACAACGCATCTCATATT3' | 60 | 73 |
| DPP7 | Forward:5' AGCCGCTGTCAGACGAGAAG 3'<br>Reverse:5' CGATCACAGCCCACCTTGAC 3' | 60 | 145 |
| RP11-23J18.1 | Forward:5' AAGCAGGCACCAAATGGAAAT 3'<br>Reverse:5' TGTGGTTGGGTTGCTGGGAT 3' | 60 | 51 |
| CEBPD | Forward:5' CTGCAGTTTCTTGGGACATAGGA 3'<br>Reverse:5' CTTCTCTCGCAGTTTAGTGGTGGT 3' | 60 | 73 |
| PDLIM1 | Forward:5' GCGAACAGCAGACCCTTAGAC 3'<br>Reverse:5' CTCCTGTTTCTCCTGAAGCATCT 3' | 60 | 96 |
| RP5-1068E13.7 | Forward:5' AGACACGCACTGGGCTTACAT 3'<br>Reverse:5' GATCAGACTGGTCTCTGGAGATTG 3' | 60 | 170 |

The relative gene expression was determined by calculating the ratio between the expression of target and housekeeping genes.

### 3. Methods for enrichment test

Functional variants have direct impact on gene expression and function. Thus, a set of functional variants with direct impact on gene expression and function are expected to be enriched in the overall disease association signals. We test enrichment in low p-values of functional variants using the VEGAS-like sum of squares test (SST)(1), which by summing the individual SNPs square statistics can be considered as a multivariate statistic. In comparison with other enrichment statistics (e.g. Simes test(2)), SST is more powerful to detect enrichment in a set of variants having many signals of small-to-moderate effect. It is well known that Simes tests can be rather conservative for SNPs in high linkage disequilibrium.

GWAS of large studies, e.g. PGC(3;4), are known to yield a fair number of statistically significant signals. Besides these significant signals, GWAS also harbor many small and moderately large signals spread over the entire genome(5). Thus, the statistics from a large GWAS, are already enriched in small p-values. Consequently, to be considered candidate to containing causal variants, a putative set of variants should be enriched for association signals “above” the background enrichment of the GWAS.

The adjustment for background enrichment of SST is as follow. Let SST statistic for the entire genome be $S = \sum_{i=1}^{k} {X_i}^2$ . It follows that $(S) = \sum_{i=1}^{k} {\mu_i}^2 + k$ , i.e. SST statistics behave like the sum of non-central $\chi_1^2$ variables having non-centrality parameter $\lambda = \frac{\sum_{i=1}^{k} {\mu_i}^2}{k}$ . Based on the scan statistics, the non-centrality pararmeter is estimated as $\hat{\lambda} = \frac{\max\{S-k,0\}}{k}$ . Consequently, under the null hypothesis of no enrichment above the GWAS background, the distribution of the square of a univariate scan statistic should be assumed to be ${X_i}^2 \sim \chi_{1,\hat{\lambda}}^2$ , not a central $\chi_1^2$ . Assume that the statistics associated with SNPs in the putative set of variants are the first $m < k$ statistics in the scan, i.e. $X_i,\ i = 1, \ldots, m$ . Then the SS statistic associated with this set of variants is $S_p = \sum_{i=1}^{m} {X_i}^2$ . If the number of SNPs is large enough for the Central Limit Theorem to provide a reasonable distributional assumption, under the null hypothesis of no enrichment above GWAS background, $S_p$ is approximately a normal variable with mean $\mu_S = m\hat{\lambda} + m$ and variance $\sigma_s^2 = (2 + 4\hat{\lambda})[m + \sum_{i \neq j} Cor(X_i^2, X_j^2)]$ , i.e. $S_p \sim N(\mu_S, \sigma_s^2)$ . To test for enrichment above background in the putative set of variants, we compute the normally distributed statistic $Z = \frac{S - m\hat{\lambda} - m}{\sigma_S}$ and test $H_0: E(Z) = 0$ vs. $H_a: E(Z) > 0$. The only unknown is $\sigma_S$ which can be estimated with reasonable accuracy from LD reference genotype data, e.g. European subjects from the 1000 Genomes (Joober, 2011) available in Mach (Li et al., 2010) and Impute (Williams et al., 2012) databases. However, most tested set of variants do not have the very large number of SNPs needed to ensure that the Central Limit Theorem provides a good approximation for the distribution of the enrichment statistic, $Z$. Consequently, we use 50,000 simulations to assess the statistical significance of $Z$ under a more restrictive "competitive" $H_0$, i.e. we test for an enrichment beyond the enrichment of the remaining SNPs in the GWAS (or the average GWAS enrichment). For these simulations we assume that i) the LD structure between selected SNPs is the one estimated from the European subjects of 1000 Genomes project and ii) the distribution of the square statistics is a non-central chi-square ( ${X_i}^2 \sim \chi_{1,\hat{\lambda}}^2$), i.e. it is not central chi-square as used in a more liberal self contained $H_0$. In more detail, similar to VEGAS (Liu et al., 2010) , the statistics associated with SNPs on a chromosome are assumed to have a multivariate normal distribution with the variance matrix equal to the correlation between SNPs' genotypes.
